# Sources of suboptimality in a minimalistic explore-exploit task

**DOI:** 10.1101/348474

**Authors:** Mingyu Song, Zahy Bnaya, Wei Ji Ma

## Abstract

Balancing exploration and exploitation is a fundamental aspect of decision-making. It remains unknown whether people are close to optimal in striking this balance, and if not, how exactly their behavior deviates from optimality. Many existing paradigms are not ideally suited to answer this question, as they contain complexities such as non-stationary environments, stochasticity under exploitation, and reward distributions that are unknown to participants. Here, we introduce a task without such complexities, in which the optimal policy is to start off exploring and to switch to exploitation at most once in each sequence of decisions. The behavior of 49 laboratory and 143 online participants deviated both qualitatively and quantitatively from the optimal policy, even when allowing for bias and decision noise. Instead, people seem to follow a suboptimal rule in which they switch from exploration to exploitation when the highest reward so far exceeds a certain threshold. Moreover, we show that this threshold decreases approximately linearly with the proportion of the sequence that remains, suggesting a novel temporal ratio law. Finally, we find evidence for “sequence-level” variability which is shared across all decisions in the same sequence. Our results provide a new perspective on the explore-exploit dilemma, and emphasize the importance of examining sequence-level strategies and their variability when studying sequential decision-making.

---

Many daily-life decisions involve a tradeoff between exploration and exploitation [6, 17] ‒ should you stick with the same brand of breakfast cereal or try a new one? Should you stay in your current job or explore new opportunities? Exploitation generally means choosing the action believed to have the maximum expected reward, while exploration is choosing any other action, which may be beneficial in the long run [22]. It is not well understood how and by how much people deviate from optimality in explore-exploit decisions. Studying such optimality is hard because the pure explore-exploit problem is often intertwined with multiple cognitive processes: the reward distributions may be unknown to participants and require learning [1, 21]; in order to evaluate options, participants need to remember and aggregate a series of past actions and rewards [8]; sometimes, the outcome of an exploitation action is also uncertain [8, 1, 21]; the environment might be non-stationary, either because it changes over time [8, 15], or because the agent’s decisions affect the environment [7]; and the number of consecutive decisions might be stochastic (indefinite decision horizon), which requires the estimation of remaining opportunities to exploit any new information [19, 1].

To investigate optimality in the absence of these complexities, we introduce a paradigm in which the participant has full information about the task structure and the only form of stochasticity is the one intrinsically associated with the outcome of an explorative action. The participant was told that they were a tourist in a foreign city. Restaurant ratings in this city ranged from 1.0 to 5.0 and followed a truncated Gaussian distribution, which was visualized on the screen throughout the experiment (**Figure 1a**). Each trial consisted of 5 to 10 virtual “days”. On each “day”, the participant decided to go either to a random new restaurant (exploration) or the best restaurant so far (exploitation), and received a reward equal to the rating of the restaurant. Their goal was to maximize the total reward of the entire trial.

We denote the trial length by *T*, the reward by *r*, the action by *a* (0 for exploitation and 1 for exploration), the highest reward received so far in a trial by *r*\*, and the number of days left by *t*_left_,. The optimal policy in this task depends only on *r*\* and *t*_left_; thus, the optimal agent does not need to maintain or update the reward distribution. For a given state (*r*\*, *t*_left_), the optimal agent compares the expected future reward *Q*(*r*\*,*t*_left_; *a*) between *a* = 0 and *a* = 1. This *Q* function is specified by the Bellman equations, which we calculate using the value iteration algorithm [3] (see Supplementary Material section 1 for the derivation of the optimal policy). The optimal agent explores more when *r*\* is lower and when there are more days left (**Figure 1b**). Since over the course of a trial, *r*\* increases and *t*_left_ decreases, the optimal policy switches from exploration to exploitation at most once in a trial (see Supplementary Material section 2 for a formal proof of this property for any deterministic reward distribution, and section 3 for the calculation of the expected number of switches under the current reward distribution).

**Figure 1:**
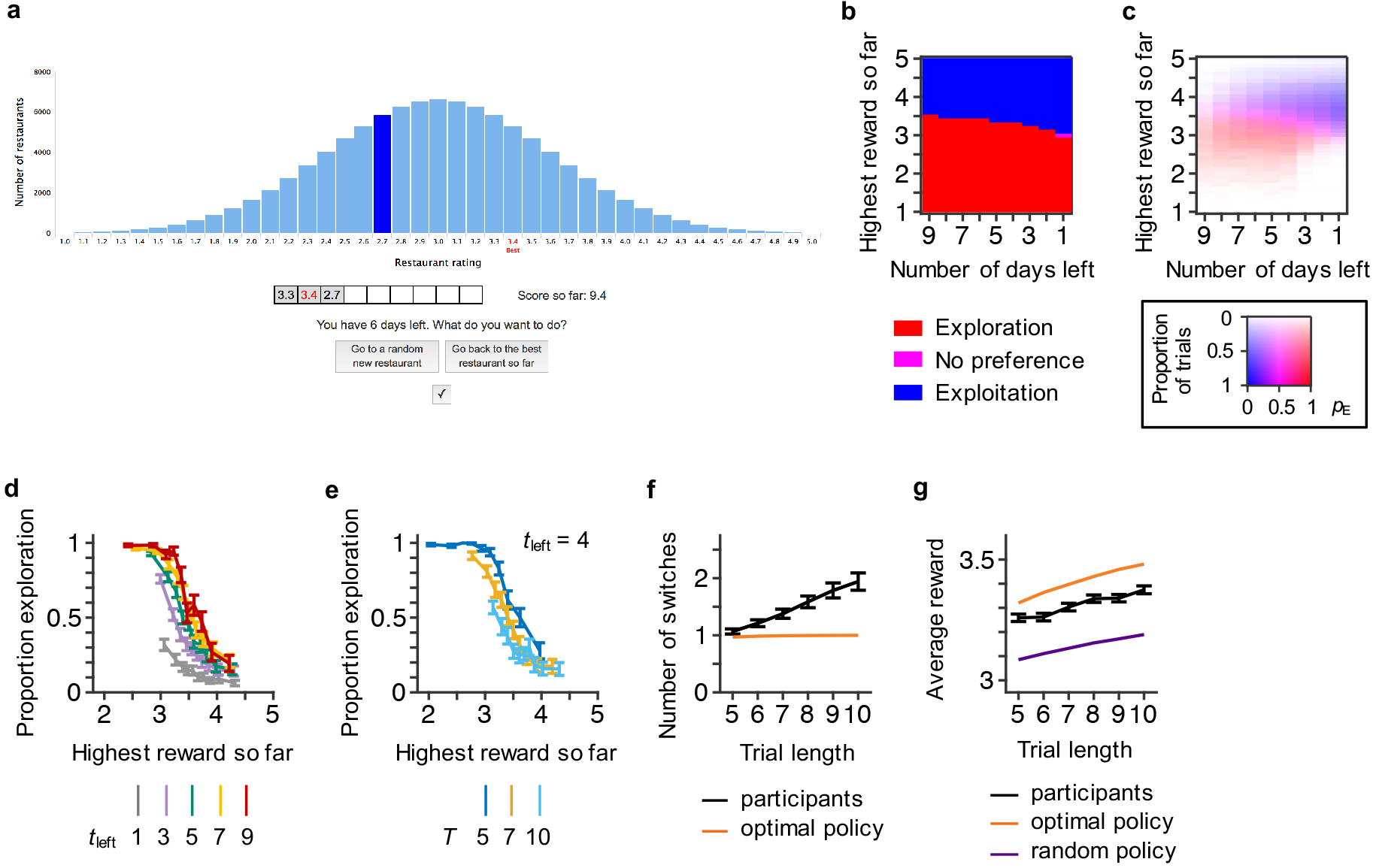
Experimental design, optimal policy, and summary statistics. **(a)** Example of a response screen. The histogram shows the distribution of restaurant ratings. Dark blue bar: most recent reward. Red text: highest reward so far. The history of rewards and the “score so far” (accumulated reward) are shown below the histogram. In this example, the participant is in the fourth day of a 9-day trial and received 2.7 on the third day; the highest reward received so far is 3.4 and the accumulated reward is 9.4. **(b)** Under the optimal policy the decision of whether to explore only depends on the highest reward so far (*r*\*) and the number of days left (*t*_left_). The optimal agent explores when *r*\* is low and t_left_ is high, **(c)-(g)** Summary statistics of data from laboratory participants. Here and elsewhere, error bars indicate 1 s.e.m. across participants, **(c)** Proportion of decisions in which participants explored as a function of the highest reward so far and the number of days left, averaged across participants. The color code is two-dimensional: hue represents the proportion of exploration (*p*_E_), and saturation = log(1 + proportion of trials)/log(2). **(d)** Slices from the plot in (*c*). For each participant and each *t*_left_, we divided the *r*\* values from all decisions into 10 quantiles; within each quantile, we calculated the proportion of decisions in which the participant explored. We plotted the mean and s.e.m. of that proportion against the mean across participants of the median *r*\* in that quantile. **(e)** Proportion of exploration as a function of the highest reward so far for *t*_left_ = 4, broken down by trial length, aka total number of days (*T*). In the optimal policy, *T* would be irrelevant, **(f)** The number of switches between exploration and exploitation, averaged across trials, as a function of trial length, for participants (black) and for the optimal policy (orange; see calculation in Supplementary Material section 3). **(g)** Average reward as a function of trial length for participants (black), the optimal policy (orange), and the random agent (purple). In (d) and (e) we only show part of the data; see Figure S1 for the full data. Panels (c) through (g) show qualitative and quantitative deviations of human behavior from the optimal policy.

We tested 49 laboratory participants and 143 online participants using Amazon Mechanical Turk (see Methods for details). All participants passed a task comprehension test. Each laboratory participant completed 180 trials (1170 choices), and each online participant completed 60 trials (390 choices). At the end, laboratory participants were asked to write about their strategy. In the main text, we report the results from laboratory participants; results and conclusions are consistent for online participants (see Supplementary Material section 5). We characterize the choice data using a set of summary statistics **(Figure 1c-g).** Participants’ choices **(Figure 1c)** resembled a stochastic version of the optimal policy **(Figure 1b).** Participants explored more in the beginning of a trial than towards the end, and when the highest reward received so far was lower **(Figure 1c-d).** Logistic regression on the choice against *r*\* and *t*_left_ returned coefficients of ‒5.64 ±0.61 (mean ± s.e.m.) for *r*\* (*t*(48) = ‒9.20,*p* < 0.001; here and elsewhere, *t*-tests are all two-tailed) and 0.545 ± 0.038 for *t*_left_, (*t*(48) = 14.4,*p* < 0.001).

**Figure 2:**
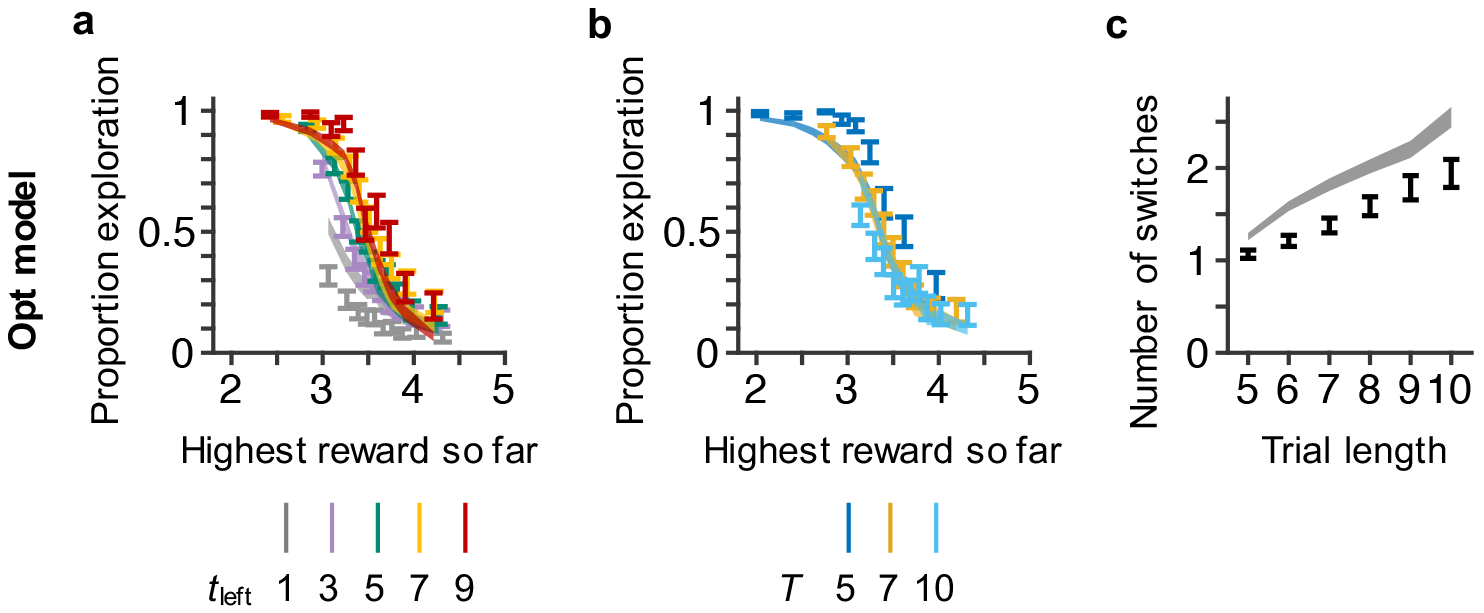
Fits of the Opt model to selected summary statistics. The Opt model fits poorly despite allowing for bias and decision noise, **(a), (b)** and **(c)** correspond to the summary statistics in Figure Id, le and If respectively. Error bars are data; shaded areas are model fits (both are 1 s.e.m).

Participants’ behavior, however, also showed clear deviations from the optimal policy. Their choices were influenced not only by *r*\* and *t*_left_, but also by the trial length *T*. For a given *t*_left_, participants explored more in shorter than in longer trials **(Figure 1e**, **S1b and S2)**. Adding *T* as a regressor in the logistic regression returned a coefficient of –0.276±0.036 for *T* (*t*(48) = −7.73,*p* < 0.001); the coefficients for *r\** and *t*_left_ remained significantly different from zero (both *p* < 0.001). Furthermore, we noted before that the optimal agent switches at most once per trial. Participants switched more frequently than optimal for any given trial length (**Figure 1f**; all *p* < 0.01 after Bonferroni correction). The number of switches also increased with trial length (linear regression slope: 0.180 ± 0.024, *t*(48) = 7.44,*p* < 0.001). Participants’ average reward per “day” was significantly higher than the average reward of an agent who randomly explores or exploits, but also significantly lower than an optimal agent (**Figure 1g**; in both comparisons, *p* < 0.001 for every trial length, after Bonferroni correction). Participants earned a higher average reward per day on longer trials (linear regression slope: 0.0242 ± 0.0029, *t*(48) = 8.36.*p* < 0.001; **Figure 1g**). Taken together, these results showed that participants adopted a reasonable but suboptimal strategy. Similar to the optimal agent, participants tended to switch to exploitation towards the end of a trial, and when the highest reward received so far was high. However, their choices were more stochastic and were influenced by trial length; on average, they switched more often between exploration and exploitation and earned less reward than the optimal agent.

Some of the observed deviations from optimality could simply be due to decision noise. Therefore, we first consider a model obtained by adding softmax decision noise [16] to the optimal policy. The probability of exploring is then

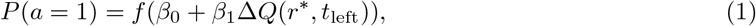

where Δ*Q*(*r*\*, *t*_left_) ≡ *Q*(*r*\*, *t*_left_ 1) − *Q*(*r*\*, *t*_left_; 0) and 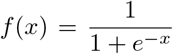 is the logistic function. When the inverse temperature *β*_1_ is higher, behavior is closer to deterministic. The parameter *β*_0_ captures a bias towards exploration or exploitation. We call this model the Opt model. We fitted the Opt model to individual choices for each individual participant using maximum-likelihood estimation. The fits to the summary statistics are qualitatively similar to the data (the three most diagnostic summary statistics are shown in **Figure 2**; see **Figure S3** for model fits to the rest summary statistics), but quantitatively, the Opt model does not fully capture the influence of number of days left on proportion of exploration (**Figure 2a**), nor could it predict the effect of trial length on choices (**Figure 2b**). The Opt model also overestimates the number of switches (**Figure 2c**). These results indicate that the Opt model is not a good description of participants’ choices, and that the deviation of human behavior from the optimal policy is not merely due to bias and softmax decision noise.

**Figure 3:**
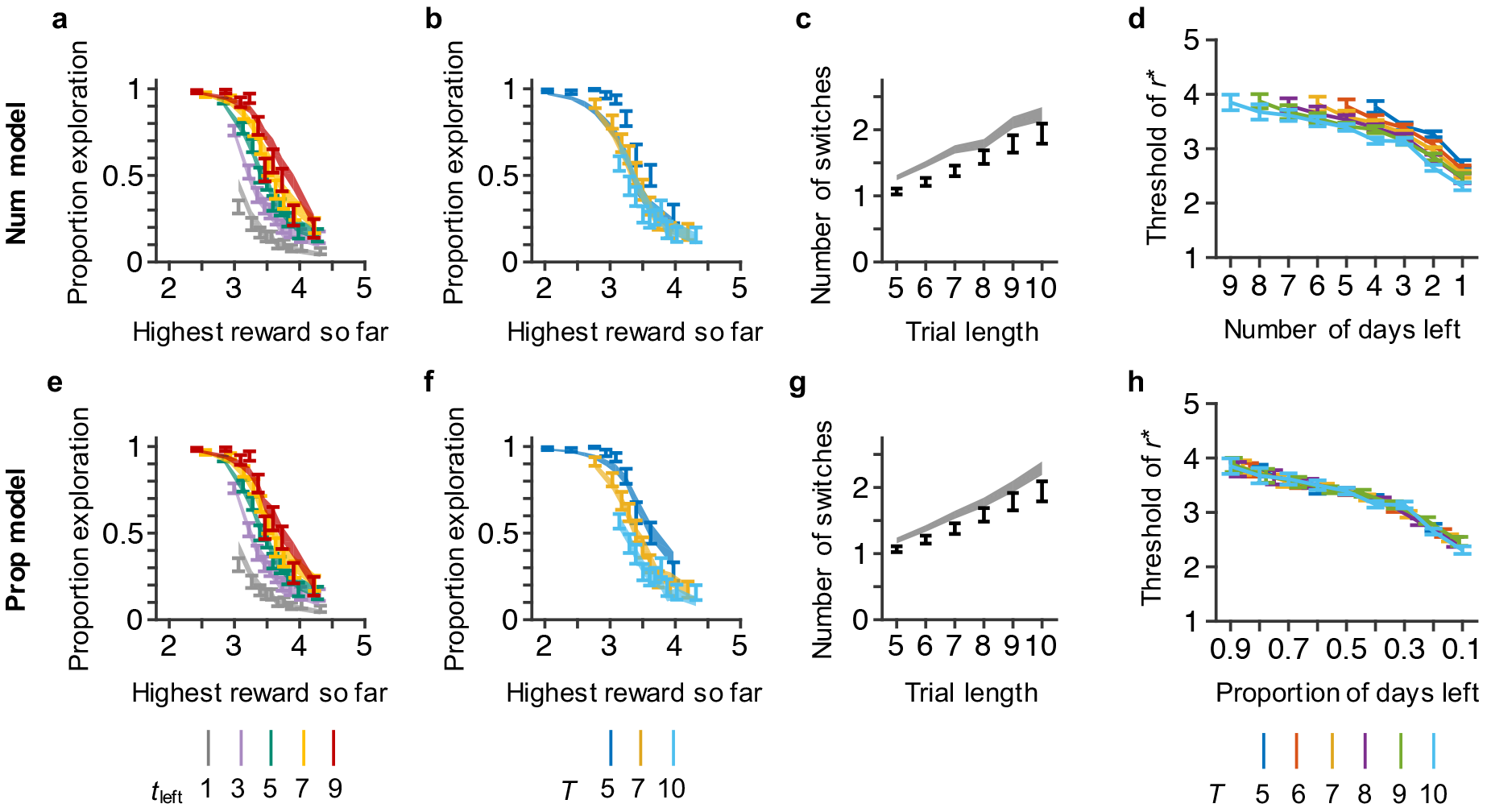
Evidence of a threshold-rule depending on the proportion of days left. Participants’ behavior is better characterized by a threshold rule that depends on the proportion of days left than by one that depends on the number of days left. This suggests that relative time is more important than absolute time, **(a, b, and c) and (e, f, and g)** Model fits of the Num and Prop models to the summary statistics in Figure 1d, 1e, and 1f. **(d)** The fitted threshold of *r*\* as a discrete function of *t*_left_ and *T*. **(h)** The same curves as in (d) with the independent variable changed to proportion of days left (each curve is stretched along the x axis respectively).

We next consider the possibility that deviations from optimality are not only due to bias and noise but also due to systematic suboptimalities in the policy. Since calculating optimal future values could be computationally demanding, people might instead use a simple heuristic [11, 20]. Indeed, participants reported using heuristic strategies (see Supplementary Material section 9): most participants (30 out of 43 valid responses) reported to have explored in the beginning of a trial and switched to exploitation when the best restaurant rating reached a threshold. Some reported that the threshold was related to trial length and where they were in a trial. Therefore, we consider threshold rules, in which the agent starts out exploring and once *r*\* exceeds a time-dependent threshold, switches to exploitation and keeps exploiting until the end of the trial. We still allow for softmax decision noise and bias (inherent in the threshold function). Denoting the threshold by *θ*, the probability of exploring is then

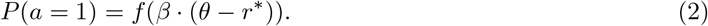

Inspired by the optimal policy, we first consider a model whose threshold function also depends on the number of days left (*t*_left_) but in a linear way: *θ* = *kt*_left_ + *b* (*k* and *b* are model parameters). We call this the Num model. The Num model fits better to the proportion of exploration as a function of *t*_left_ (**Figure 3a**). However, as can be expected, the dependency of participants’ choice on trial length (*T*) would not be predicted (**Figure 3b**), because there is no information about *T* in the Num model.

To explore the specific threshold strategy people used, we fit a Flexible-Threshold model where *θ*(*t*_left_,*T*) is a discrete function of *t*_left_ and *T* (**Figure 3d**; 40 free parameters, including 39 thresholds and a softmax noise). The fitted threshold depends both on *t*_left_ and *T*: it decreases over the course of a trial, and is lower on longer trials. When we change the independent variable from the number of days left (*t*_left_) to the proportion of days left (*t*_left_/*T*), the thresholds of different trial lengths overlap with each other, indicating that participants used a threshold rule that almost linearly depended on the proportion of days left (**Figure 3h**). Thus, we consider a model where *θ* = *kt*_left_/*T* + *b*, and call it the Prop model. The Prop model provides a better fit to the data than the Num model or the Opt model (**Figure 3e-g**). Specifically, it fits well to the effect of *T* on proportion of exploration (**Figure 3f**); it also preserves the good fit on the influence of number of days left (**Figure 3e**). These findings suggest that the relevant quantity in deciding when to switch to exploitation is the relative rather than the absolute time in the trial. This perhaps reflects an organism’s need to plan across multiple different time scales [14].

**Figure 4:**
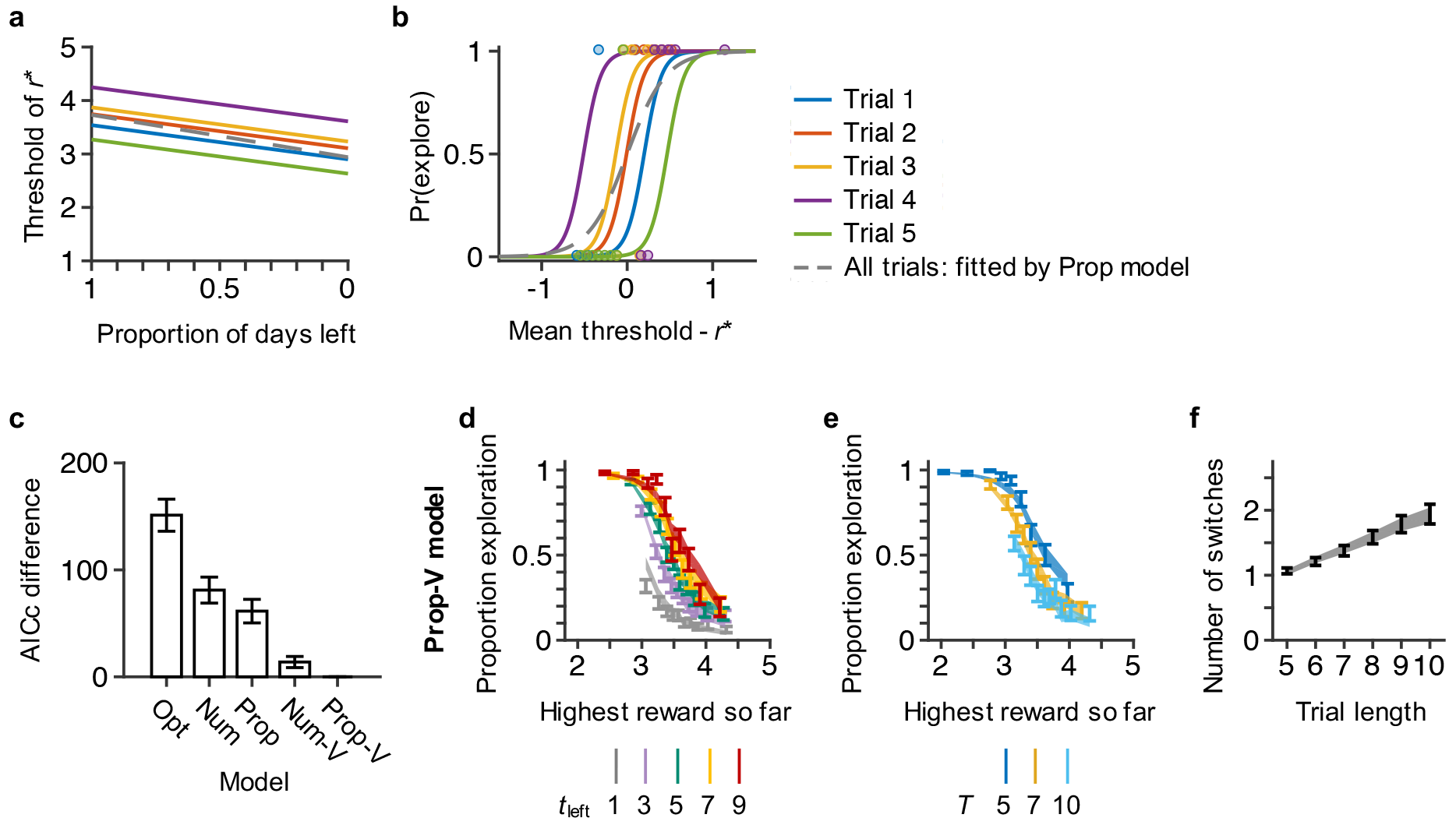
Sequence-level variability as implemented through the variable-threshold models (Num-V and Prop-V). The models with sequence-level variability account better for participants’ choices, **(a)** We postulate that choices are not only subject to softmax noise, but also to variability at the level of an entire sequence of choices (a trial), as implemented through random trial-to-trial shifts in the threshold function (five example trials shown). Here we use the Prop-V model as an example, but the same idea applies to the Num-V model. **(b)** Corresponding random shifts in the psychometric curves of the probability of exploration as a function of the difference between mean threshold and *r*\*. Dots are data simulated using Prop-V model for the five example trials; the simulated data are then fitted using the Prop model and the resulting psychometric curve is shown in dashed grey. If Prop-V (Num-V) is the true model, fitting the Prop (Num) model will overestimate the choice-level softmax noise (the dashed grey line is shallower than the colored lines), and therefore the number of switches per trial, **(c)** Comparison of all five models. The AICc values of the the Prop-V model are used as baseline. Both variable-threshold models fit better than their fixed-threshold counterparts. **(d), (e), and (f)** Model fits of the Prop-V model to the summary statistics in Figure 1d, 1e, and 1f.

All the models we have considered so far systematically overestimate the number of switches (**Figure 2c**, **Figure 3c** **and 3g**), which could result from the overestimation of softmax decision noise. This motivates us to introduce another type of variability: a random shift of thresholds at the level of an entire trial, which consists of a sequence of 5 to 10 choices (**Figure 4a**). Different from choice-level softmax noise, which changes the slope of the psychometric curve, sequence-level variability randomly shifts the psychometric curve horizontally on each trial (**Figure 4b**). If an agent has variable thresholds but we fit their data using a fixed-threshold model (Num or Prop), the choice-level softmax noise would be overestimated to account for the sequence-level variability, and as a result the number of switches would be overestimated.

We implement sequence-level variability as a Gaussian random variable *η* with a mean of 0 and a variance of *σ*^2^. Thus the probability of exploration for the variable-threshold models will be

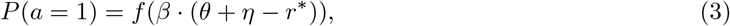

where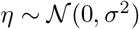. We add this threshold variability to the Prop and Num models to obtain the Prop-V and Num-V models, respectively. This addition greatly improves model fits in both cases (**Figure 4c**, AICc decreased by 61 ± 11 from Prop to Prop-V, and by 67 ± 11 from Num to Num-V), and both models fit well to the number of switches (**Figure 4f** and **Figure S4c**). The good fits to other summary statistics are preserved for the Prop-V model (**Figure 4d** **and 4e**). The Num-V model still could not fit to the effect of trial length on the proportion of exploration (**Figure S4b**).

We next examine the possibility that the Prop-V model fits better than the Prop model not because of sequence-level variability but because the linear threshold assumption is incorrect and the model tries to compensate for that. We compare the Prop-V model to the Flexible-Threshold model without sequence-level variability (**Figure 3h**), which can account for threshold functions of arbitrary shapes. The Prop-V model fits much better (AICc difference: 156 ± 34), indicating that the good fit of the Prop-V model cannot be attributed solely to threshold mismatch.

Next, we ask whether learning can account for sequence-level threshold variability. Fitting the Prop-V model separately to the first and second halves of the task shows no difference in the parameter estimates (*p* > 0.27 for all four parameters). Moreover, the average reward per trial was not different between the first and second halves of the task (*p* = 0.76). Therefore, we consider it unlikely that sequence-level threshold variability is due to learning.

To sum up, we designed a simplified explore-exploit task that allowed us to quantitatively characterize the deviations of human behavior from optimality. We found that human participants behaved qualitatively similar to the optimal policy, but that they adopted a heuristic decision rule instead of calculating optimal values. In particular, they scaled their decision threshold to the trial length. This suggests a temporal scaling law for planning that might be easier and more intuitive to implement than using the absolute amount of time left. It is broadly consistent with the notion that ratios of temporal durations, not absolute values, are relevant in interval timing [10] and long-term memory [5]; however, it is not clear whether the similarity is more than superficial.

In addition to choice-level softmax noise as commonly used to model value-based decisions, participants also showed sequence-level variability. Even though we only considered a very simple form of such variability, it underlines the point that when studying sequential decisions, one has to examine policies and their variability at the level of the entire sequence rather than only at the level of individual decisions.

Methodologically, the sources of suboptimality that we found would have been hard to identify with a more traditional design, as they might have been confounded with failures of learning or memorization. The minimalistic features of our task ‒ in particular the ability to perform an exploitation action without much stochasticity, and a fixed, known horizon ‒ are not only a modeling convenience, but potentially also approximate some real-world decisions: the problem was originally motivated by author W.J.M. having to decide whether to go to the same breakfast place in Beijing during his few days of visit there. Another example could be choosing a brand of diapers of a certain size, knowing that the baby will grow out of that size within a month. However, even when a real-world scenario contains more types of uncertainty than we included, our approach might still be useful as a starting point for process models that dissect sources of suboptimality.

A similar simplified task was used by Sang and colleagues [18]. Participants were asked to maximize the total score across a sequence of 20 decisions, each of which consisted of either flipping a card from a deck of 100 cards, numbered from 1 to 100, or picking one of the cards they had already flipped. Like in the present study, participants explored more when the highest number so far was lower and when there were more decisions left in the trial; people also switched more between exploration and exploitation than the optimal policy. However, no other summary statistics were analyzed and potential sources of suboptimality were not modeled; in particular, because only one trial length was tested, it was not possible to compare threshold rules.

The current study has several limitations. First, in the threshold models, we assumed that the threshold depends linearly on the decision variable, but other forms could be explored. Second, although we showed participants the reward distribution throughout the experiment and tested their understanding of the distribution, they might perceive probabilities and/or rewards in a distorted way [13], resulting in a distorted reward distribution (We have tested this by adding in a risk-attitude parameter to the Prop-V model; it does not improve the fit much: AICc difference −4.1 ± 1.6; **Figure S5**). Participants might have a changing belief of the distribution depending on the rewards they received, which could be the origin of the sequence-level threshold variability. This is an open and interesting question to explore, and could be pursued by future studies.

Our work also provides a new angle in the search for the neural basis of explore-exploit behavior. Previous work has found involvement of rostrolateral prefrontal cortex in exploration actions[8, 4, 2]. Our paradigm could potentially be useful to examine to what extent this activation can be broken down into “pure” exploration (as studied here) versus learning about the reward distributions. Moreover, dorsolateral prefrontal cortex (dlPFC) has been found to encode task-relevant variables for decision making[9]. Thus, we would expect to find signals correlated with the decision threshold and/or the decision variable (difference between the threshold and the highest reward so far) in dlPFC; we would also expect to find sequence-level variability in the corresponding fMRI signal.

## Methods

### Experiment 1

#### Task

Participants were placed in a computerized task environment with a backstory in which they were a tourist in a foreign city. A trial consisted of *T* “days” (*T* varies pseudo-randomly between 5 and 10, with equal number of each). On each of the *T* days, they had to decide where to go for dinner. The rating of the restaurants in the city varied between 1.0 (worst) to 5.0 (best); the distribution was a renormalized truncated Gaussian with mean 3.0 and standard deviation 0.6. The number of restaurants was assumed to be much greater than 10. The histogram of restaurant’s rating was shown on the screen. On each “day” except for the first one, the participant would choose to click one of two buttons ‘Go to a random new restaurant’ or “Go to the best restaurant so far”; on the first “day”, only the first button could be clicked. After clicking on either button, a check mark button would appear in the bottom center of the screen, and the button that did not get chosen would be disabled (not shown in Figure 1a). Participants needed to click on the check mark button to confirm their choice. After a choice was confirmed, the resulting rating would be shown (e.g. “2.9”) and the corresponding bar would be highlighted in the distribution. If the exploration button was clicked, the rating would be drawn randomly from the Gaussian distribution. If the exploitation button was clicked, the rating would be the highest rating obtained so far in the trial. It was shown on the screen the number of “days” left, the highest rating so far (marked in red), the numerical history of rewards (with the highest one in the series highlighted), and the score (sum of ratings) so far in the trial. At the end of the trial, the participant was told their trial score and the next trial commenced. The task was self-paced with no time limit.

#### Procedure

Participants were explained the task and the distribution of possible rewards using a series of self-paced instruction screens. To test their understanding of the distribution, they were shown the histogram with a simple two-alternative question of the type “If you randomly go to a restaurant in this city, which rating is more probable? A. 2.5 B. 4.5” (the correct answer would be A). Participants had to correctly answer at least two out of three such questions to be allowed to continue; participants who did not meet this criterion would be sent home. The remaining participants were shown the example screen in Figure 1a with the different elements annotated to familiarize them with it. After the instruction phase, participants completed 180 trials in a single session, each consisting of 5 to 10 decisions. After completing all 180 trials, we asked laboratory participants “what kind of strategy did you use in the task?”, to which they typed an answer (no length limits). See Supplementary Material section 9 for full answers to the question. At the end of the session, we randomly chose one trial and determined a bonus payout based on the score. Participant were told this procedure, and we emphasized to them that each trial was equally important in determining their payout.

#### Participants

49 participants (10 male, 24 female and 15 unknown, aged 18-57), participated in the experiment. All of them passed the qualifying test as described in *Procedure*. Participants received $10 for completing the experiment (about 45 minutes), plus a performance bonus up to $5. The experiments were approved by the University Committee on Activities Involving Human Participants of New York University. Each participant gave informed consent before the experiment.

### Experiment 2: Online experiment

Experiment 2 was conducted on Amazon Mechanical Turk, programmed using the psiTurk interface [12]. We obtained separate IRB approval for this project from the University Committee on Activities Involving Human Participants of New York University. Each participant gave informed consent before accepting the task. The procedure was the same as in Experiment 1, except for the following differences: (a) Participants completed only 60 trials. (b) Participants were paid nothing if they did not pass the qualifying test, $1.5 if they passed the test and completed the experiment, and a performance bonus of up to $5. (c) Participants were recruited through the general Amazon Mechanical Turk task listing. (d) Due to technical problems, participants’ reported strategies were not recorded. 143 participants participated (no demographic information provided).

### Experiment 3, 4 and 5

Experiment 3 was the same as Experiment 2, except that most history information (trial length, previous rewards and the accumulated reward so far) was hidden from participants (**Figure S8**). 131 participants participated (no demographic information provided).

Experiments 4 and 5 were conducted before other three experiments. They were the same as experiments 1 and 2 except that we did not require participants to click on a confirm button after each choice. The old design was thought to bias participants towards continuing the same choice. However, results turned out to be consistent with or without the confirm button. 16 (7 male and 9 female, aged 20-43) and 108 (no demographic information provided) participants participated in experiment 4 and 5 respectively.

Results from experiments 3, 4 and 5 were consistent with the main experiments 1 and 2 (**Figure S9-S14**), so we take them as replicates of the main experiments.

### Models and model fitting

#### Constrains on thresholds

For all threshold models (including the Num, Prop, Num-V and Prop-V models), we didn’t set constrains on parameters *k*, *b* and *σ*, so the threshold *θ* is unbounded. In order for the thresholds to be cognitively meaningful, we forced *θ* to be between 1 and 5 by adding the following constrain after each calculation of threshold:

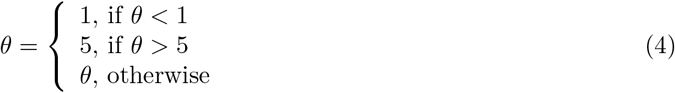

#### Algorithm for optimization

For the Opt model, we used mnrfit function in Matlab. For Num, Prop, Num-V and Prop-V models, we used fminunc function in Matlab for maximum likelihood estimate, and we ran each fitting for 100 times from random starting points to get the global maximum.

#### Synthetic data testing

Synthetic data test was conducted for all the models and model parameters were recovered correctly.

#### Fits to the summary statistics

The fits to some summary statistics (Figure 1f and 1g) were done by running the model forward using the fitted parameters, generating simulation data from it, and extracting summary statistics in same way as for real data.

### Data availability

All data in this paper are available at https://github.com/mingyus/explore-exploit (under folder data/).

### Code availability

All codes used in this paper are available at https://github.com/mingyus/explore-exploit.

## Author contributions

All authors designed the study, developed the models, and wrote the paper. M.S., Z.B. collected the data and performed the analyses.

